# Intranasal GHK peptide enhances resilience to cognitive decline in aging mice

**DOI:** 10.1101/2023.11.16.567423

**Authors:** Matthew Tucker, Addison Keely, Joo Young Park, Manuela Rosenfeld, Jackson Wezeman, Ruby Mangalindan, Dan Ratner, Warren Ladiges

**Author notes:** **Correspondence.** W Ladiges.

## Abstract

Brain aging and cognitive decline are aspects of growing old. Age-related cognitive impairment entails the early stages of cognitive decline, and is extremely common, affecting millions of older people. Investigation into early cognitive decline as a treatable condition is relevant to a wide range of cognitive impairment conditions, since mild age-related neuropathology increases risk for more severe neuropathology and dementia associated with Alzheimer’s Disease. Recent studies suggest that the naturally occurring peptide GHK (glycyl-L-histidyl-L-lysine) in its Cu-bound form, has the potential to treat cognitive decline associated with aging. In order to test this concept, male and female C57BL/6 mice, 20 months of age, were given intranasal GHK-Cu, 15 mg/kg daily, for two months. Results showed that mice treated with intranasal GHK-Cu had an enhanced level of cognitive performance in spatial memory and learning navigation tasks, and expressed decreased neuroinflammatory and axonal damage markers compared to mice treated with intranasal saline. These observations suggest that GHK-Cu can enhance resilience to brain aging, and has translational implications for further testing in both preclinical and clinical studies using an atomizer device for intranasal delivery.

## Introduction

The ability to respond to and recover from a physically stressful event is defined as physical resilience [1]. Based on the geroscience concept that aging and age-related disease conditions are associated with the same pathways [2,3], resilience to aging, as measured by ability to recover from physical stress, provides a resistance environment for preventing or delaying the onset of age-related conditions. Brain aging and cognitive decline are aspects of growing old [4]. Age-related cognitive dysfunction ranges from memory lapses associated with typical aging to early stages of neurodegenerative conditions. Dementia is the term used for more severe cognitive impairment where patients experience memory loss as well as difficulties in maintaining independence in normal living activities [5]. Mild cognitive impairment is often used to describe the stage between cognitive decline related to aging and dementia [6]. Age-related cognitive impairment (ARCI) entails the early stages of cognitive decline, and is extremely common, affecting millions of older people.

The brain, similar to other organs, ages as the result of alterations in multiple molecular pathways referred to as hallmarks of aging [7]. Different organs age at different rates depending on the aging pathways affected [8]. Factors involved in brain aging are not yet fully understood, but using anti-aging drugs to target one or more aging pathways is a rational approach to treating cognitive dysfunction [9]. Investigation into ARCI as a treatable condition is relevant to a wide range of cognitive impairment conditions, since mild age-related neuropathology increases risk for more severe neuropathology associated with MCI and dementia associated with Alzheimer’s Disease [5].

Mouse models are highly effective in age related studies as the shortened timeline allows observation of the full impact of treatments in months as opposed to the years of human clinical trials. ARCI occurs in C57BL/6 mice and is characterized by reductions in learning ability, working memory, and other focus-based assessments [10]. These mice at older ages are an excellent model to test drugs to understand and develop effective therapeutic approaches for more severe forms of cognitive impairment including dementia. A recent publication showing that the peptide glycyl-L-histidyl-L-lysine (GHK) prevented sleep-deprived learning impairment in aging C57BL/6 mice [11] suggested that GHK has the potential to treat cognitive dysfunction associated with aging.

GHK is a naturally occurring tripeptide released from the parent protein SPARC during proteolytic breakdown [12]. In the event of an injury, GHK supports angiogenesis, remodeling, and tissue repair as a copper complex (GHK-Cu). The peptide is clinically approved as a topical application for age-related skin conditions, and promoted mainly as a skin rejuvenation drug [13]. Using the Connectivity Map gene profiling software developed by the Broad Institute, a search found GHK, out of several thousand biological molecules, could alleviate the impairment of TGFβ1 [14] shown to be associated with neurodegeneration [15]. In addition, GHK has been shown to be an endogenous antioxidant by decreasing hydroxyl and peroxyl radicals [16], and improving cognitive performance in aging mice [11]. Unpublished observations in our lab suggest that GHK-Cu complex is more effective than unbound GHK while unbound Cu has no effect in several in vitro and in vivo experiments.

GHK is a peptide and not orally bioavailable. Since there is little knowledge of how efficiently the peptide crosses the blood brain barrier when administered parenterally, an intranasal delivery route was selected for this study. Intranasal delivery of peptides to the brain relies primarily on extracellular passive transport from the olfactory region of the nasal cavity [17, 18, 19]. The lattice formed by overlapping epithelial cells creates a pathway for molecules to traverse from the mucosal layer of the nasal cavity to the lamina propria [20, 21]. Size restrictions for these junctions between the epithelial cells fluctuate but the high turnover of cells creates a window of increased permeability that can be leveraged by a translocating neurotherapeutic [22]. Passive diffusion from the lamina propria can then occur through the perineural space that exists between non-neuronal cells that encapsulate the axons to the brain [20, 23, 24], allowing direct access to neurons.

We show in this study that aging C57BL/6 mice given intranasal GHK-Cu for two months were more resilient to developing brain aging phenotypes than mice treated with intranasal saline.

## Methods

### Animals and experimental design

C57BL/6Jnia male and female mice, 20 months of age, were obtained from the National Institute on Aging Aged Rodent Colony, contracted by Charles River, Inc. Mice were housed in a specific pathogen free (SPF) mouse facility at the University of Washington main campus in Seattle, WA. An SPF mouse facility is defined as a clean space free of common mouse pathogens and verified by viral and bacterial tests (IDEXX Bioanalytics). The status of the room was monitored under the guidance of the Rodent Health Monitoring Program within the purview of the Department of Comparative Medicine at the University of Washington. The detection of infectious agents not acceptable in UW rodent SPF housing was performed in concert with similar standards at Charles River and the NIA aging rodent colony. All mice utilized in the study were group housed, up to five per cage, and given nestlets (Ancare Corp, Bellmore, NY) to provide activity and stimulation. Mice were monitored for health issues daily, and cages were replaced biweekly.

Mice were conditioned to handling procedures for two weeks to reduce stress response and acclimate the mice to intranasal drug delivery [25]. One week after the conditioning procedure, treatment was started and continued daily for two months, when mice were tested for behavioral performance, euthanized and brains collected for neuropathology. All procedures were approved by the University of Washington IACUC.

### GHK-Cu dose and Delivery

GHK was used as a GHK-Cu complex (Active Peptide, Cambridge, MA) at a dose of 15mg/kg. Cu ions can induce toxicity in mice if dosage exceeds levels of 35 mg/kg [26]. Within the GHK-Cu complex, Cu made up 14% of the total molecular content, so a 15mg/kg GHK-Cu dose had 2.1 mg/kg copper. Mice were given 15 mg/kg in 20 ^µ^ L saline daily between 7 AM and 10 AM over an 8-week period.

The study utilized a custom 3D printed nasal atomizer designed with Fusion360 software. The device consisted of 1) a fitted pressure opening for attaching to a standard syringe, 2) a thin and flexible double layered plastic tube, and 3) a hard plastic concentric ring tip (Figure 1). The device worked by pressure from a syringe plunger, which pushed the fluid down the tube and past the concentric ring tip where the flow was converted into a thin mist that exited the applicator at the entrance to one nostril directly into the nasal cavity. The device was 3D printed out of a 504nm photopolymerizable resin with soybean oil as a binder. It was designed with a smooth interior surface contoured towards the tip to reduce fluid loss in the tip to a negligible volume. The radius of the tip was 1.25mm, small enough to enter the nostril of a mouse, but the width increased by approximately a millimeter down the tip length as a natural stop to prevent entering too deep into the nasal cavity. This feature provided a standard depth for fluid discharge. The tip measured 2.17 mm in length from face to contact, so it was long enough to reach the target epithelium with the fluid discharge without irritating or damaging the nasal cavity with the applicator length, and short enough to not be easily broken off or damaged. The ring design ensured a relatively even distribution of the atomized droplets on the proximal and distal regions of the superior nasal cavity. The base of the device included three layers of ridges to interact with the surface of a universal Luer Lock syringe with a shortened tip. Two ridges fit snugly around the diameter of the syringe tip and provide a pressure seal to prevent fluid loss. The third ridge was flush with the base of the device itself. The use of a pressure fit syringe in contrast to a standard Luer Lock threaded syringe allowed for faster transition between tip exchanges.

**Figure 1.**
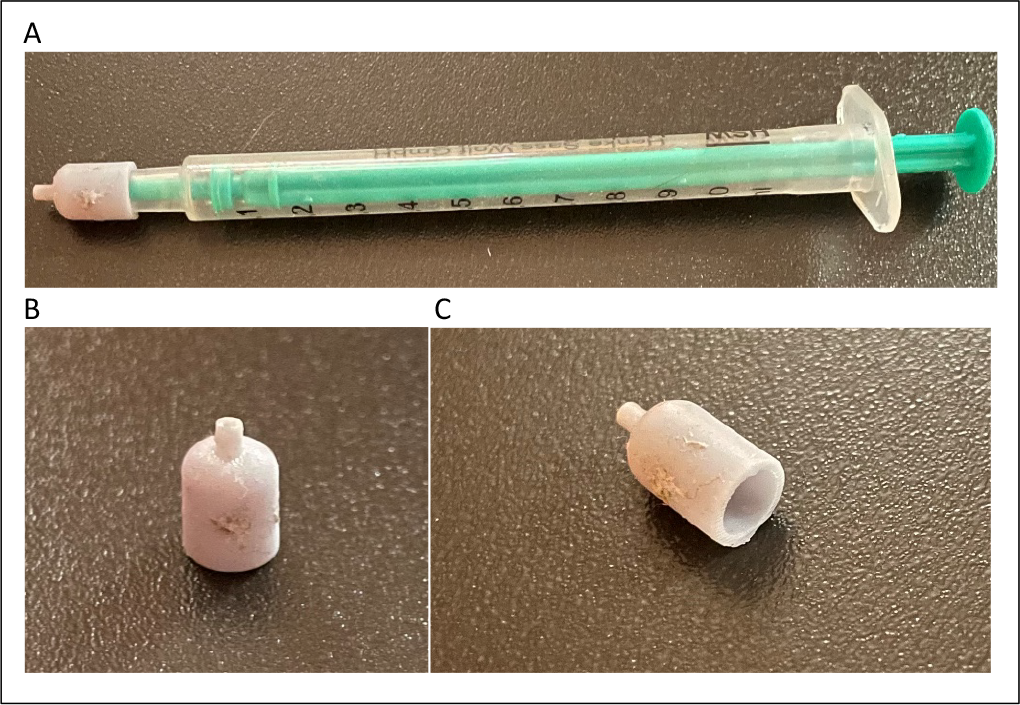
**A**. 3D printed intranasal atomizer tip affixed to a 1mL pressure syringe **B**. Atomizer vertical view. **C**. Atomizer isometric view.

### Behavioral tests

#### Y-maze memory task

Cognitive function was assessed with the Y-maze, a behavioral spatial test of working memory [27]. The Y-maze consists of three arms of equal length spaced 120° apart, with raised walls along each arm. Each mouse roams the maze for five minutes while their path through the three arms is tracked by entry into each arm. The test is designed to measure spontaneous alternation, which is the capacity of a mouse to naturally explore an arm of the maze instead of returning to an arm they have more recently visited, as an indicator of working memory over a short period of time. Data was generated from each trial over time when the test was administered in weeks 0, 4, and 8 of the study. Data were recorded as a percentage of the number of times the mouse completed a triad or loop through all three arms throughout the series of arm locations during the trial time divided by the number of entries minus two. This created the percentage of spontaneous alternation for each mouse each time the test was run, with increased alternation percent indicative of increased memory performance [28]. Additional data from the number of arm entries were also collected.

#### Spatial navigation learning task

Mice were also tested using a spatial navigation learning task designated as the Box Maze [29]. Mice were introduced to a large foil-lined square plastic box with the stressor of a bright overhead light. Each wall of the box possessed two floor-level holes for a total of 8 holes, with seven blocked by an inset cap and one left open connected to an s-curved escape tube to a darkened empty mouse cage. The escape hole set-up gives the illusion that each hole leads into darkness so the mouse will need to learn where the escape hole is in order to escape more quickly upon repeated trials. Four consecutive trials were given to each mouse with a time limit of 180 seconds. Between each trial there was a 30 second resting period. Data were recorded from each trial as the time, in seconds, it took each mouse to find and fully enter the escape hole. If a mouse did not enter the escape hole within the time limit of 180 seconds, a time of 180 seconds was recorded.

#### Neuropathology assessment

Mice were euthanized with CO_2_ inhalation followed by decapitation for brain segmentation. Brain sections from one hemisphere were separated into frontal cortex, hippocampus and cerebellum, quick frozen in liquid nitrogen and stored at −80°C. Brain sections from the other hemisphere were fixed in 10% buffered formalin for 48 hours before being blocked in paraffin wax and cut into 5um slices onto glass slides for immunohistochemistry (IHC).

IHC staining was performed as previously described [30]. Slides were rehydrated with xylene, decreasing concentrations of ethanol, and finally deionized water. Slides were incubated in a 1:10 Citrate Buffer Antigen Retrieval solution at 98°C in a hot water bath for 20 minutes for antigen retrieval. Staining was performed using an AbCam IHC kit according to manufacturer instructions. The slides were incubated with primary antibodies in a Tris-buffered saline, 0.1% Tween® 20 Detergent solution (TBST) overnight in a humidified chamber at 4°C. Slides were incubated with Biotinylated Goat Anti-Mouse and Streptavidin Peroxidase with TBST washes in between. Finally, DAB chromogen was applied to the slides and incubated followed by washes with TBST and deionized water. Slides were then dehydrated with increasing ethanol concentrations and xylene before being mounted with mounting media and a coverslip.

The antibodies used were specific for monocyte chemoattractant protein-1 (MCP-1) as an indicator of inflammation, ionized calcium-binding adaptors (IBA-1) as a marker for microglia, and neurofilament light-chain-1 (NFL-1) as indicator of axonal damage. IHC stains were photographed using a microscopic camera at 4-to-20X scale with images in each brain region for each mouse. Each photo was processed through QuPath digital imaging as previously described to create a color gradient heat map to quantify the stain density standardized to control stains [31].

#### Statistical Analysis

The statistical analysis of data was determined between the control and variable cohorts. Quality independent variables such as sex were also included as per cohort standards. Mice were all housed and received the same treatment protocol and handling procedures, differing on just the drug (GHK-Cu), and were assessed with the same behavioral and histological tests and indicators. Differences between variables were assessed via a two-tailed t-test with scientific significance at p≤ 0.05.

## Results

### Mice treated with intranasal GHK-Cu showed enhanced cognitive performance

Male and female mice, 22 months of age and given intranasal GHK-Cu for 8 weeks, had a higher spontaneous alternation percentage than intranasal saline cohorts (Figure 2A) indicating increased cognitive ability. Interestingly, after 4 weeks of treatment, only males experienced higher alternation percentages indicating females required a longer period of time to respond to treatment. Arm entry data showed that males, but not females, treated with intranasal GHK-Cu had a lower number of entries per trial than mice treated with intranasal saline at both 4 weeks and 8 weeks (Figure 2B) indicating a sex dependent effect of the peptide.

**Figure 2.**
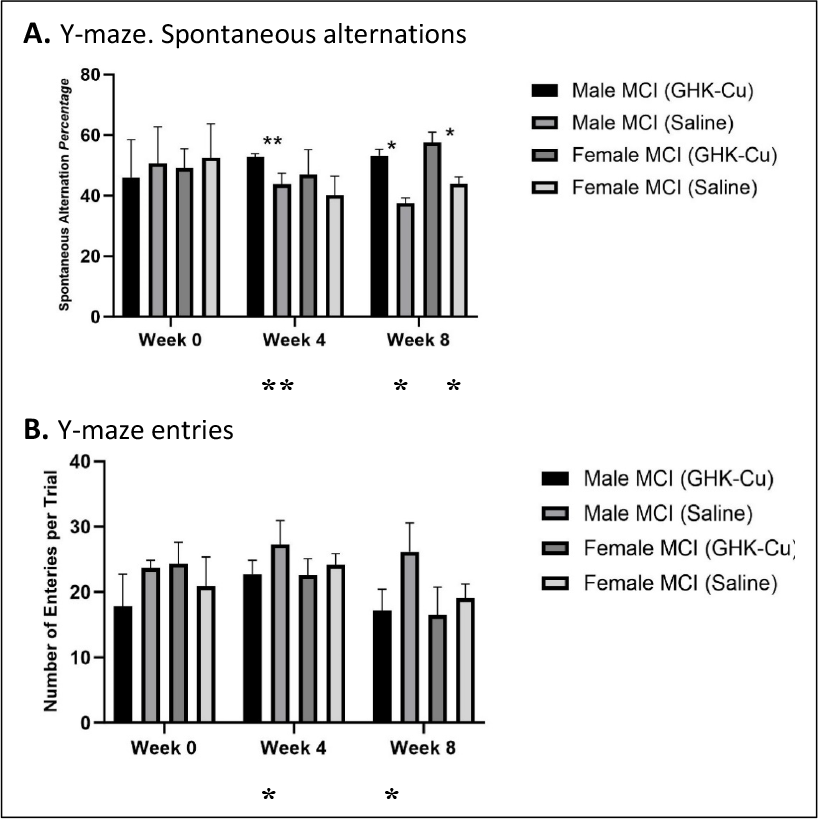
Y-Maze behavioral test results. **A**. Both males and females treated with intranasal GHK-Cu showed increased spontaneous alternation at week 8 compared to mice treated with intranasal saline. **B**. Males, but not females, treated with intranasal GHK-Cu showed a decreased number of arm entries at weeks 4 and 8 compared to males treated with intranasal saline. **p≤0.05, *p≤0.01. N = 17-19/cohort. MCI = mild cognitive impairment/age-related cognitive impairment.

Male and female mice treated with intranasal GHK-Cu showed decreased escape times by the fourth trial in the Box Maze compared to mice treated with saline (Figure 3A). Both male and female mice treated with GHK demonstrated larger magnitudes of improvement over the four trials compared to male and female given intranasal saline, as both a comparison over each trial, and as a total sum of improvement (Figure 3B). Male and female GHK-Cu cohorts and saline cohorts had comparable values for escape hole attempts on the first trial, but male and female mice given intranasal GHK-Cu demonstrated fewer escape hole attempts on their final attempt to escape compared to the intranasal saline cohorts (Figure 3C).

**Figure 3.**
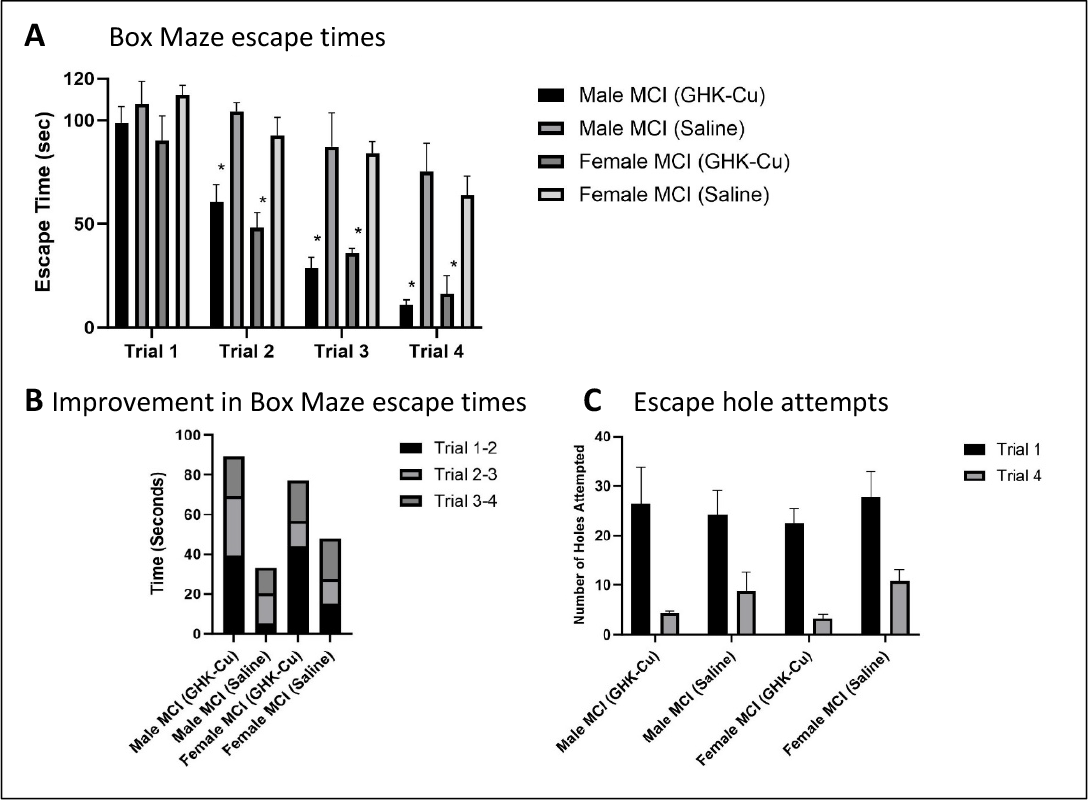
Box maze escape data. A. Both male and female mice treated with GHK-Cu found the escape hole more quickly than mice treated with intranasal saline. **B**. The magnitude of escape times was greater in trials 2-4 in mice treated with GHK-Cu compared to mice treated with intranasal saline. **C**. Mice treated with intranasal GHK-Cu had fewer escape hole attempts until success compared to mice treated with intranasal saline. *p≤0.01. N = 17-19/cohort. MCI = mild cognitive impairment/age-related cognitive impairment.

### Mice treated with intranasal GHK-Cu showed attenuation of brain aging parameters

MCP-1 levels in female mice treated with intranasal GHK-Cu were significantly lower in the frontal cortex compared to intranasal saline treated female mice indicating a decrease in neuroinflammation, while males only showed a nonsignificant trend of lower MCP-1 levels (Figure 4A). Intranasal GHK-Cu treatment had a nonsignificant suppressive effect on microglia in males but not females as demonstrated by staining intensities of IBA-1 (Figure 4B). Both male and female mice treated with intranasal GHK-Cu had significantly lower concentrations of NFL-1 in the frontal cortex compared to intranasal saline treated cohorts (Figure 4C) indicating GHK-Cu was able to attenuate axonal damage associated with brain aging in C57BL/6 mice.

**Figure 4.**
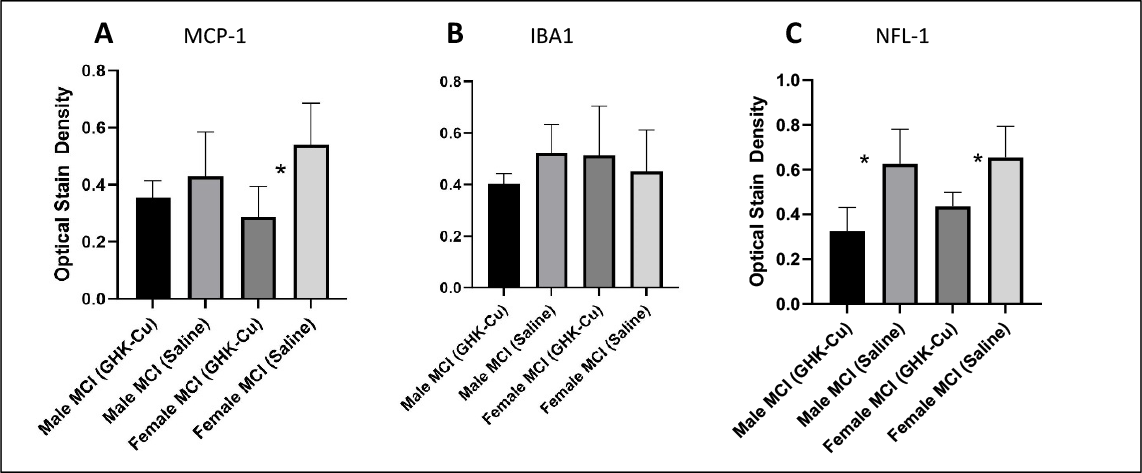
Immunohistochemistry staining intensities in the frontal cortex generated by Qu-Path digital imaging. **A**. MCP-1. **B**. IBA-1. **C**. NFL-1. *p≤0.05. N = 9-12/cohort. MCI = mild cognitive impairment/age-related cognitive Impairment.

## Discussion

GHK-Cu was tested in this study as an intranasal neurotherapeutic for aging C57BL/6 mice that naturally show cognitive decline with increasing age. Spontaneous alternation percentage, an indication of working spatial memory in the Y-Maze test, increased in male but not female mice after 4 weeks of treatment with intranasal GHK-Cu. However, after 8 weeks of treatment with intranasal GHK-Cu, when mice were 22 months of age, both males and females showed an increase in alternation percentage. This indicated that mice treated with GHK-Cu had an enhanced level of cognitive performance compared to mice treated with intranasal saline. In addition, there was a lower number of arm entries for males, but not females, treated with intranasal GHK-Cu compared to mice treated with intranasal saline after weeks 4 and 8, which supports the alternation percentage findings, at least for males. Even though there was a trend for improved cognition in females, the peptide was not as effective as for males over the same treatment period. A longer treatment period for females might result in more effective results.

Mice treated with intranasal GHK-Cu were able to recall the correct escape hole from visual landmarks in the Box Maze test, and tried 8-10 times fewer escape holes during the final trial compared to their first trial. Learning latency was significantly pronounced in both male and female mice treated with intranasal saline, with much slower escape times and a lower magnitude of improvement, validating the naturally occurring cognitive decline of these mice. In addition, saline treated cohorts experienced 4-6 fewer escape hole attempts from their first to their final trial. Therefore, the combined complementary results of the Y-Maze test and the Box Maze test show that 20-month-old C57BL/6 mice treated with intranasal GHK-Cu for 8 weeks have increased resilience to cognitive decline compared to control mice treated with intranasal saline.

The neuropathology data supported the cognitive test results. Inflammation is a hallmark pathway of aging and is increasingly present in the brains of C57BL/6 mice [32]. The brains from mice treated with intranasal GHK-Cu showed a decrease in MCP-1, a marker used to indicate inflammation [33]. Interestingly, the decrease in staining intensity was more pronounced in the brains from female mice than male mice, even though the peptide showed a more effective increase in resilience to cognitive decline in males. The lack of significant differences in staining intensity for IBA-1, a marker for microglia, was somewhat surprising. Microglia are resident immune cells in the brain and are generally increased with increasing age as the result of neuronal insults and damage [34]. Brains from male mice but not female mice, treated with intranasal GHK-Cu, only showed a decreased trend for the presence of microglia, suggesting microglia are not a robust target for GHK-Cu at least under the experimental conditions of this study. The third IHC neuropathology antibody marker used in this study showed an increased staining intensity for neurofilament light chain (NFL-1) as an indicator of axonal damage [35]. Clearly, the brains from both male and female mice treated with intranasal GHK-Cu showed decreased staining intensity for NFL-1 suggesting that axonal damage is associated with brain aging that can be attenuated with GHK-Cu treatment. Although the mechanism for this effect was not investigated, it could be speculated that GHK-Cu might be triggering a wound healing effect similar to that occurring in damaged skin [13]. Overall, these neuropathology observations suggest that GHK-Cu acts on molecular markers associated with different pathways in the brain in a sex dependent manner, and that additional drugs could be added as a cocktail to be more effective in enhancing resilience to age-related cognitive decline.

The intranasal atomizer device has a number of advantages in delivering peptide drugs and other small molecules directly to the brain of mice. GHK-Cu was efficiently administered and effective in suppressing the age-related cognitive decline in mice in this study. The use of the device in mice required a 2-week acclimation period in order to efficiently deliver GHK-Cu into the nasal cavity without excessive and stressful restraint. Preliminary training was also necessary to properly use the device under light restraint conditions in fully awake mice without the need for any analgesics or anesthetics.

In conclusion, intranasal GHK-Cu was shown to be effective at abating cognitive decline in aging C57BL/6 mice over an 8-week treatment period. This observation suggests that GHK-Cu can enhance resilience to brain aging, and has translational implications for further testing in both preclinical and clinical studies using an atomizer device for intranasal delivery.

## Acknowledgments

This work was supported by National Institutes of Health grants R01 AG057381 and R01 AG057381 (Ladiges PI).

